# BMP-2 Release from Biomaterial Carriers: Reporting Heterogeneity and the Limits of Cross-Study Prediction

**DOI:** 10.64898/2026.11.13.766607

**Authors:** Warren Zhou, Leon Zhou, Wanda Hou, Yunzhi Peter Yang

## Abstract

Published release profiles offer a potential basis for predicting bone morphogenetic protein-2 (BMP-2) delivery from biomaterial carriers, but heterogeneous reporting and study-specific effects complicate their reuse in machine learning (ML) models. As a preliminary exploration, we compiled 79 release experiments and 487 time points from 21 publications with a workflow that combined a large language model assistant and human verification, and we examined data comparability and cross-study prediction with ML models. Only 38 experiments (48%) reported BMP-2 alone as a percentage of the loaded dose. Within individual studies, binding components and coating layers were associated with slower release, consistent with established carrier mechanisms. Using carrier family as the sole predictor, a family-median model and gradient-boosted decision trees achieved lower mean absolute errors than a global-median baseline under random hold-out validation (0.42 and 0.44 versus 0.52 log_10_ units). Publication-grouped cross-validation did not establish a corresponding improvement (0.57 and 0.48 versus 0.50). Only 10 of the 28 modeling experiments belonged to carrier families represented in the training data under this scheme, and carrier family was strongly associated with publication. These findings highlight the need to distinguish within-study interpolation from cross-study generalization. Broader coverage of each carrier family, quantitative formulation and assay descriptors, and accessible numerical source data are needed to support predictive modeling of BMP-2 release.

## 1 Introduction

Recombinant human bone morphogenetic protein-2 (rhBMP-2) is an osteoinductive growth factor used clinically for bone repair. Its performance depends in part on the kinetics of protein release from the delivery carrier. The approved absorbable collagen sponge [1] provides limited retention: approximately 50%–80% of the loaded BMP-2 is released within 24 hours, with approximately 5% remaining after two weeks [2]. Clinical administration consequently uses milligram-scale doses, although BMP-2 is biologically active at nanogram levels. A critical review of industry-sponsored spinal fusion trials estimated adverse event rates of 10%–50%, depending on surgical approach, approximately 10–50 times the rates reported in the original publications [3].

Carriers developed to control BMP-2 release include ceramics, synthetic polymers, hydrogels, polysac-charides, and natural proteins. BMP-2 contains a heparin-binding site [4]; affinity-based carriers, including heparin-conjugated materials, exploit this interaction to prolong retention [5,6]. However, release profiles are typically presented as figures in individual publications, with substantial variation in reporting units and assay conditions. This heterogeneity limits quantitative comparisons across studies and the use of published data to predict release from new formulations.

Machine learning (ML) has been applied to literature-derived datasets in related areas of drug delivery. Bannigan et al. compiled 181 release profiles comprising 3783 fractional release measurements for 43 drug–polymer combinations of polymeric long-acting injectables [7]. Among eleven algorithms, LightGBM achieved a mean absolute error (MAE) of 0.114 for fractional release in nested cross-validation grouped by drug–polymer combination. Bao et al. released a dataset of 321 in vitro release experiments and 4913 time points for drug-loaded poly(lactide-co-glycolide) microparticles, extracted from 113 publications [8]. He et al. curated 714 assay records for nanoparticle drug delivery systems and compared eleven algorithms for classifying biological activity using three-fold cross-validation [9]. These studies paired published measurements with formulation descriptors. Bannigan et al. and Bao et al. digitized release figures using plot-digitizing software; Bao et al. additionally used two independent reviewers for screening and data collection.

Reliable reuse of published measurements requires explicit provenance and verification. Large language models (LLMs) can assist extraction, but their outputs require assessment against the primary sources. In a study within systematic reviews, LLM-assisted extraction with human verification achieved 91.0% accuracy across 9341 data elements, compared with 89.0% for human-only extraction [10]. A separate evaluation found that chance agreement accounted for part of GPT-4’s apparent screening and extraction accuracy [11]. Source references and independent checks are therefore essential for distinguishing documented measurements from unsupported values and for assessing the reliability of extracted records.

To our knowledge, a cross-carrier analysis of literature-derived BMP-2 release data has not been reported. Earlier neural-network work on protein and peptide release used 68 microparticle formulations, as summarized by Bannigan et al. [7]. BMP-2 presents additional challenges because release depends on protein– carrier binding, while the literature spans diverse materials, assays, and measurement conventions.

Here, we examine a literature-derived BMP-2 release compilation as an exploratory case study of data comparability and validation design. Specifically, we characterize the extraction workflow and assembled records, evaluate the accuracy and repeatability of figure-derived measurements, identify records that share a payload and reporting scale, and test whether carrier family predicts half-release time for publications excluded from training. We further identify the data and reporting practices needed to support more reliable cross-study prediction.

## 2 Materials and Methods

### 2.1 Overview of the Workflow

The workflow separates candidate identification, deterministic processing, and source verification (Figure 1; Table 1). An LLM assistant proposed candidate publications and records; scripts imported, normalized, and checked the data; and human readers digitized figures, verified measurements, and adjudicated inclusion and labels. Release and outcome values were accepted only if traceable to numerical source data, a table, a text statement, a figure, or an explicit calculation from reported values. Model-generated text was not treated as a measurement source.

**Table 1.** Responsibilities for candidate identification, deterministic processing, and human verification at each work-flow stage.

| Stage | LLM assistant | Scripts | Human reader |
|---|---|---|---|
| Paper selection | Ranked papers by likely relevance and marked papers for full-text review | Imported the paper list, built the screening lists, and fixed the corpus at the cutoff date | Decided on inclusion and exclusion |
| Data extraction | Proposed candidate release curves, table values, outcome endpoints, and possible problems | Required identifiers, payload, provenance, source reference, and notes for every record | Digitized figures, copied values from tables and text, and resolved unclear curve assignments |
| Source data | Identified the relevant source data sheets | Imported the source data and blocked replacement by values of lower provenance | Confirmed that each sheet matched the intended figure |
| Normalization and labeling | Proposed cohort assignments | Converted time to days, calculated release metrics, and assigned unit, cohort, and profile-quality labels | Approved the rules and checked the carrier family labels |
| Validation | Suggested profiles for checking | Compared image-derived values with source data and ran consistency checks | Read selected digitized points a second time |
| Outcome records | Identified outcome endpoints in each paper | Linked outcomes to release experiments by identifier | Decided on outcome categories and removed uncertain links |

**Figure 1.**
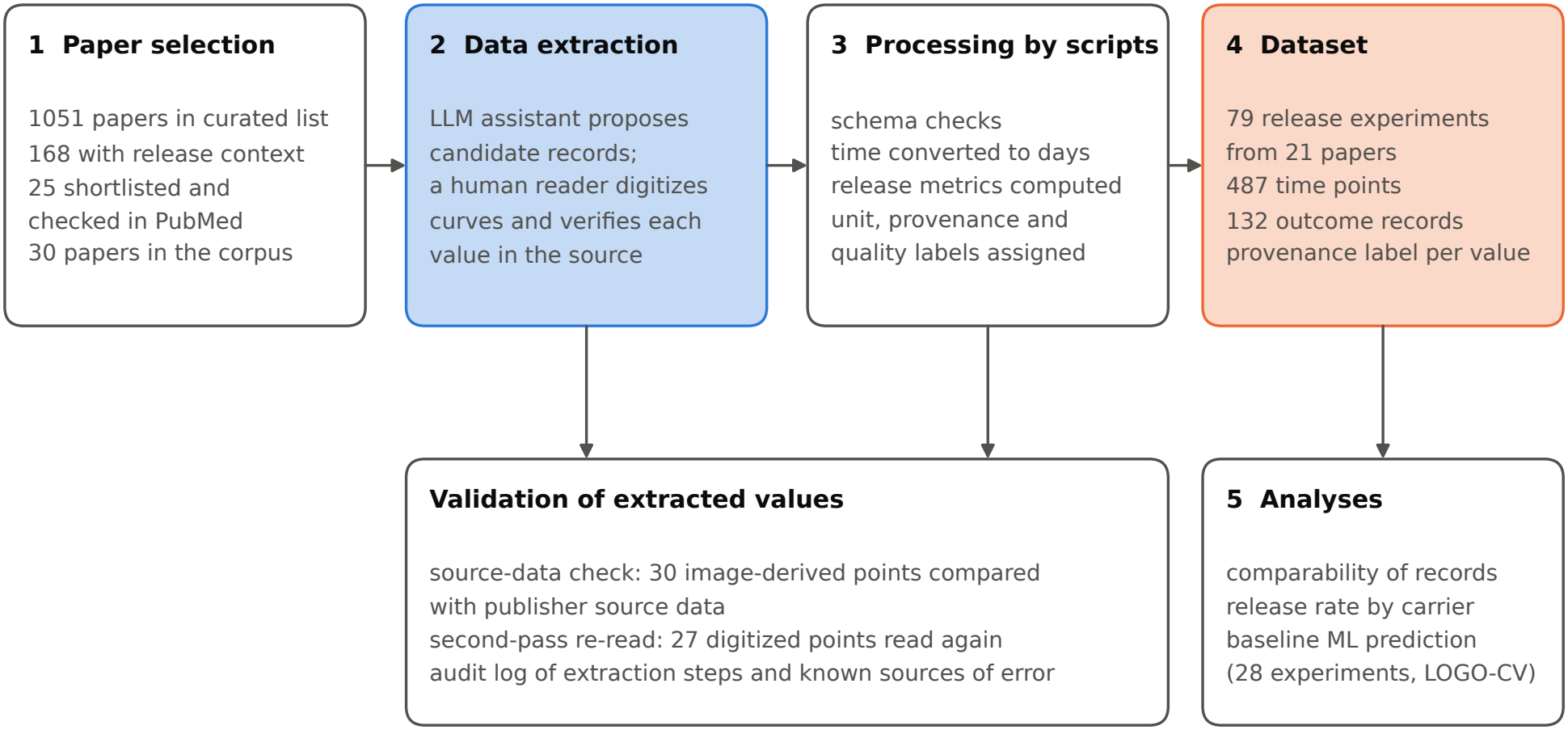
Workflow used to compile and analyze the BMP-2 release dataset. The LLM assistant proposed candidate records, a human reader extracted and verified the values, and scripts normalized and labeled the records. LOGO-CV, leave-one-group-out cross-validation; ML, machine learning.

Throughout this manuscript, *provenance* denotes the origin of a value, such as a published figure or source data file. The *payload* is the released agent. A *branch record* concerns a payload other than BMP-2 protein alone, including co-delivery with another active factor. A *cohort* groups experiments by payload type and reporting unit.

The analysis is reported through the methods, aggregate tables, and figures included in this manuscript. Its primary objective is to assess data comparability and cross-study validation in an exploratory setting.

### 2.2 Literature Source and Paper Selection

Publications were drawn from a curated spreadsheet of 1051 BMP-2 delivery papers (Supplementary Data 1) containing first author, year, title, carrier material, release context, and PubMed identifier. Records were imported programmatically. Because the spreadsheet did not originate from a systematic database search, the resulting dataset is a convenience sample rather than a comprehensive survey.

Selection was documented at each stage (Figure 1). Of the 1051 records, 168 specified an in vitro or in vivo release context. With LLM assistance, 25 were prioritized for manual screening, and their metadata, payloads, and reported release and outcome data were checked in PubMed. Fifteen were selected for initial structured extraction. Two reference papers chosen at project inception were added, yielding 17 papers. Additional papers were included as full texts, figures, or supplementary files became available, including branch studies and comparisons of in vitro and in vivo measurements. The corpus was frozen on 29 May 2026 at 30 papers. Subsequently identified papers were logged separately and excluded from the present analysis.

Selection criteria are summarized in Table 2. Core records were eligible for release-rate analyses. Branch records were retained but excluded from these analyses because a different payload or an additional active factor alters the interpretation of release and biological outcomes.

**Table 2.** Criteria for paper selection.

| **Class** | **Criteria** |
|---|---|
| Core | All of the following: exogenous BMP-2 or rhBMP-2 protein delivered locally from a biomaterial; a bone, cartilage, dental, or osteochondral regeneration context; release, retention, or elution data at defined time points; enough material detail to classify the carrier; and at least one biological outcome or bioactivity measurement. |
| Branch | BMP-2-derived peptides; plasmid, mRNA, viral, or cell-mediated expression; BMP-2 delivered together with VEGF, PDGF-BB, dexamethasone, antibiotics, or another active factor; papers on a material or platform without direct BMP-2 release data. |
| Excluded | Retracted articles and retraction notices; no measurement of payload release or retention; BMP-2 present only as an endogenous marker; time information insufficient to describe the release profile. |

A separate sample of 84 open-access papers from the same spreadsheet was used only to characterize reporting units (Section 3.3); one also belonged to the main corpus. A full-text keyword classifier identified these papers as reporting both BMP-2 release and an in vivo bone outcome. The LLM assistant inspected each full text and release figure and classified the release-axis unit as percent, mass, concentration, mixed, or unclear. These labels were not verified by a human reader and are the only unverified model-assigned labels used in this study.

### 2.3 Data Extraction and Verification

For each corpus paper, the LLM assistant examined the available full text, figures, tables, and supplementary files to propose candidate release curves, tabulated measurements, and biological outcomes. Candidate records flagged potential ambiguities, including unclear units, additional active factors, and retention measurements. A human reader then copied numerical values from tables and text or digitized curves as described in Section 2.4. Publisher source data, available for one paper [12], were imported programmatically. Acceptance and rejection rates for candidate proposals were not recorded.

Each value was stored with paper and experiment identifiers, payload, source reference, provenance label, and notes. The source reference identified the figure, table, text passage, or source data sheet; records lacking a reference were rejected. Four provenance categories were used: source data (numerical data supplied by authors or publishers), digitized figure, text, and derived. For derived values, the calculation was documented in the notes. Numerical source data were never overwritten by lower-provenance estimates.

Scripts converted times to days, separated percentage measurements from mass and concentration measurements, calculated release metrics (Section 2.6), and assigned the labels described in Section 2.5. Integrity checks verified table readability, outcome-to-experiment links or explicit unlinked status, and preservation of source data values. For the records that support the main results, an audit log recorded each extraction step, who performed and verified it, and the known sources of error.

### 2.4 Figure Digitization and Its Validation

A human reader digitized release curves by visually reading plotted points against the axes, without plotdigitizing software. Most estimates were therefore recorded as whole percentages. Before extraction, the reader documented axis quantities, units, ranges, logarithmic scaling, the measured quantity (cumulative release, retained fraction, mass, or concentration), and error-bar definitions. Points were read at published sampling times, and cumulative profiles were checked for monotonicity. For retained-fraction measurements, release was calculated as 100% minus the retained percentage, and records retained a retention label. Eleven figure-check notes documented axis interpretation and limitations such as crowded or low-resolution plots.

Two complementary checks assessed figure-derived values. For the source-data check, a script calibrated axes and detected series-specific markers by color in a published figure [12]. The resulting image-derived estimates were compared with numerical publisher data for the same panels. This check assesses image-based extraction from a clear figure rather than the accuracy of manual readings. For the second-pass check, selected manually digitized points were read again and compared with the initial readings. This check assesses repeatability; it was neither blinded nor performed by an independent second reader.

### 2.5 Dataset Structure and Labels

A release experiment denotes one formulation evaluated under one assay condition. Table 3 summarizes the dataset fields; the labels governing joint analysis are defined below.

**Table 3.**
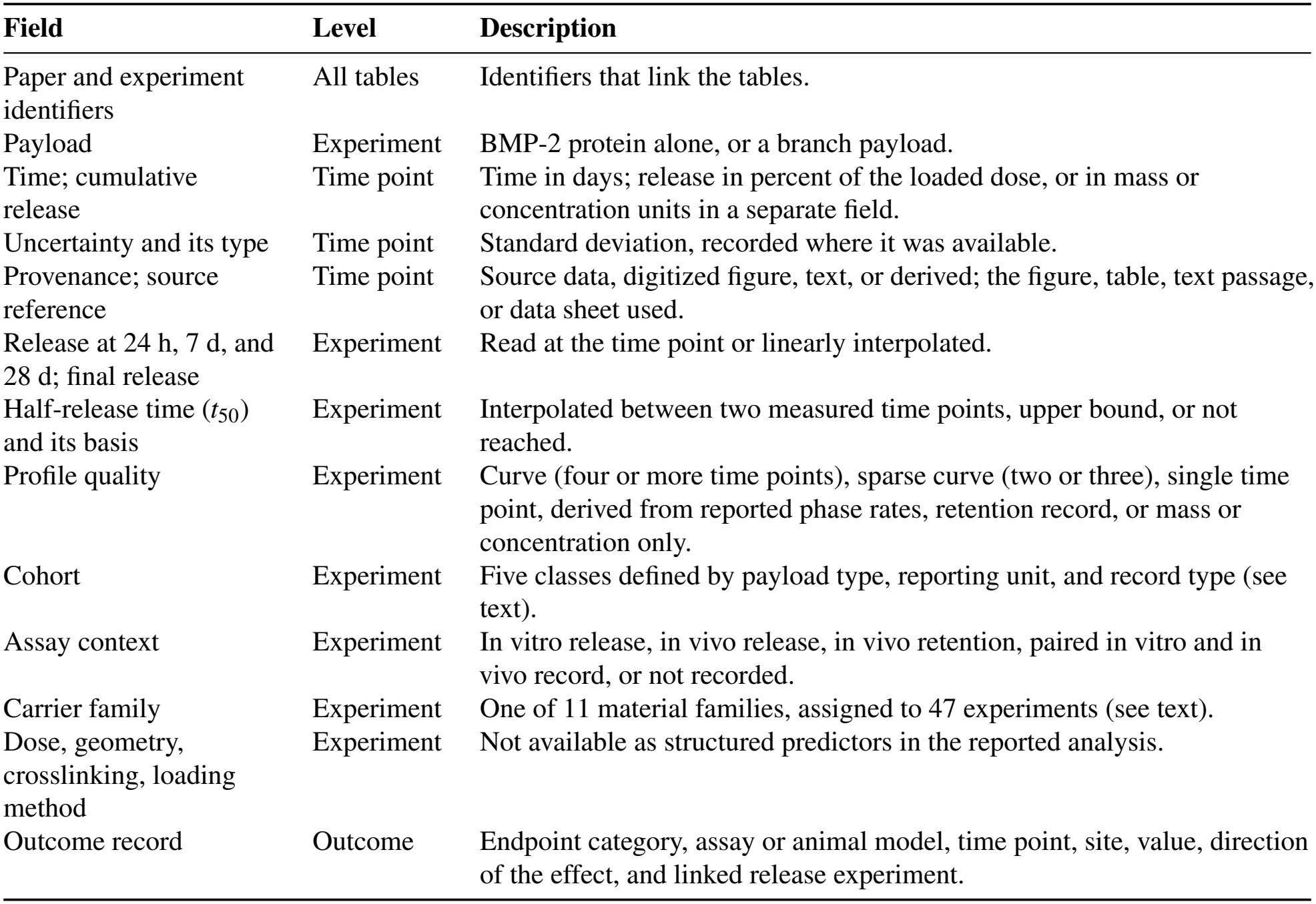
Overview of the main fields of the dataset.

Two corrections preceded analysis. First, two duplicate records were removed because the same series appeared in multiple panels of a single paper. Second, three polymer-scaffold controls without gelatin microparticles had inherited the test formulation’s family label; these were reassigned to the polymeric scaffold family. Their influence on model performance is evaluated in Section 3.5.

#### Reporting unit

Release was recorded either as a percentage of the loaded dose or in mass or concentration units. Conversion to a common percentage scale requires the loaded dose and, for concentrations, the corresponding volume and sampling information.

#### Cohort

Rule-based assignment defined five cohorts. Two comprised BMP-2-only experiments reported as a percentage of the loaded dose: one with publisher source data and one with figure-derived, text-derived, or calculated values. These *percent-release cohorts* share payload and reporting scale but differ in assay context. The remaining cohorts comprised BMP-2-only experiments reported in mass or concentration units, branch payloads (Table 2), and retention or paired in vitro/in vivo records.

#### Profile quality

Profiles with at least four time points were labeled as curves. Separate labels identified sparse curves (two or three points), single-point measurements, profiles reconstructed from reported phase rates, retention records, and mass-or concentration-only records (Table 3).

#### Assay context

Assay context was not captured as a dedicated field during extraction. It was assigned retrospectively using rules based on retention labels and experiment identifiers. Records that could not be classified this way were labeled *not recorded*.

#### Carrier family

Families represent material classes such as polymeric scaffolds, gelatin microparticles, and heparin-affinity carriers. Forty-seven experiments, including all percent-release experiments, were assigned to 11 families. The remaining 32 branch or retention records retained record-type labels without a carrier-family assignment. An explicit scripted mapping encoded classifications based on the published material descriptions. The mapping was prepared with LLM assistance and checked by a human reader, without independent classification by a second reader.

### 2.6 Release Metrics

Release at 24 hours, 7 days, and 28 days was obtained from measurements at those times or by linear interpolation between adjacent observations. Final release denotes the value at the last measured time point. Half-release time (*t*_50_) denotes the time at which cumulative release reaches 50% of the loaded dose. It was estimated by linear interpolation, assuming zero release at time zero. Profiles that did not reach 50% had no defined *t*_50_ and were excluded from release-rate analyses.

For profiles already exceeding 50% at the first observation, *t*_50_ was interpolated between the origin and that observation. For a single-point profile exceeding 50%, the observation time itself was recorded. In both cases, the first observation provides an upper bound on the true *t*_50_; these records are referred to as *upper-bound cases*. The interpolated estimate should be distinguished from that observation-time bound. No kinetic model, such as the Higuchi or Korsmeyer–Peppas model, was fitted because most measurements were digitized or derived and many profiles were sparsely sampled.

### 2.7 Machine Learning Models and Validation

The regression target was log_10_ *t*_50_, with *t*_50_ expressed in days. The modeling set comprised 28 experiments from 11 publications and 10 carrier families in the percent-release cohorts with a defined *t*_50_. Carrier family was the sole predictor and was one-hot encoded as 10 binary indicators. Although the dataset schema included dose, geometry, crosslinking, and loading method, these fields were unpopulated; no continuous carrier descriptors were available.

Three models were compared. Model complexity was restricted because the small sample size increases susceptibility to overfitting [7]. The global-median baseline predicted the training-set median log_10_ *t*_50_ for every test experiment. The family-median model used the median among training experiments in the same family, reverting to the global training median for unseen families. The third model used gradient-boosted decision trees (GBDT), implemented with scikit-learn’s HistGradientBoostingRegressor. Settings were squared-error loss, maximum depth 3, 200 boosting iterations, learning rate 0.05, and a minimum of 3 samples per leaf. Hyperparameters were fixed in advance without tuning because the sample size did not support a separate tuning set. With one categorical predictor, the model assigns identical predictions to experiments with identical encoded families. Minimum leaf size further constrains its ability to resolve sparsely represented categories. This model was included for comparison with prior tree-based release-prediction studies [7].

Two validation schemes were evaluated. In leave-one-group-out cross-validation (LOGO-CV), publications defined the groups. All experiments from one publication formed the test set, and the remaining publications formed the training set; this procedure was repeated for all 11 publications. This design evaluates generalization to an unseen study and prevents within-publication overlap between training and test sets. Such overlap can produce optimistic estimates of cross-study performance because experiments share laboratory procedures, assays, and often dose. Bannigan et al. likewise used grouped validation, with drug– polymer combination defining the groups [7]. In repeated random hold-out validation, 22 experiments were assigned to training and 6 to testing (approximately 80%/20%), irrespective of publication. This split was repeated 200 times.

Performance was quantified by MAE in log_10_ *t*_50_. Errors of 0.30 and 0.50 log units correspond to geometric mean multiplicative discrepancies of approximately 2 and 3, respectively. LOGO-CV MAE was computed across all held-out predictions and separately for experiments whose families were represented in training. Uncertainty in the MAE difference relative to the baseline was estimated by resampling the 11 publications with replacement 10,000 times. Random hold-out results are reported as the mean and standard deviation of MAE over 200 repetitions. Both schemes additionally tracked test experiments belonging to families absent from training.

Sensitivity analyses repeated both schemes for curve-quality profiles, profiles with *t*_50_ bracketed by measured time points, and experiments not recorded as in vivo. Analyses were also repeated using the corpus before the duplicate-record and family-label corrections described in Section 2.5.

### 2.8 Statistical Analysis

Group release metrics are summarized as medians. Associations of log_10_ *t*_50_ with carrier family and publication were quantified using eta squared (*η*^2^) from one-way analysis of variance. This statistic measures the fraction of total variance attributable to between-group differences. Because *η*^2^ tends to increase with the number of groups in small samples, adjusted epsilon squared (*ε*^2^) and permutation *p* values based on 10,000 permutations are also reported. The association between the two categorical variables, carrier family and publication, was measured using small-sample bias-corrected Cramér’s *V* [13], which ranges from 0 to 1. Linked biological outcomes were grouped by endpoint category, assay or animal model, time point, and anatomical site, and records in each combination were counted. Analyses were performed in Python 3.13 with NumPy 2.5 and scikit-learn 1.9, and figures were prepared with Matplotlib 3.11. The analysis scripts, input tables, and result files are provided as Supplementary Data 2.

## 3 Results

### 3.1 Dataset Overview

The corpus contained 30 publications (Supplementary Table 1), of which 21 contributed release experiments. Of the nine without extracted release experiments, five were branch papers without extracted release records and four were background or excluded records. One retracted article was retained solely to document exclusion. After duplicate removal, the dataset comprised 79 experiments (Supplementary Tables 2 and 3).

The release table contained 487 time points (Supplementary Table 4), including 443 expressed as a percentage of the loaded dose. By provenance, 352 (72%) were digitized, 78 (16%) were derived, 42 (9%) were publisher source data, and 15 (3%) were reported in text. Of 308 digitized percentage values, 253 (82%) were whole numbers.

**Table 4.** Half-release time (*t*_50_) by carrier family in the modeling set (28 experiments). Upper-bound cases exceed 50% release at the first observation. Entries reproduce the analysis estimates, including origin-based interpolation where used. A dagger marks families containing upper-bound cases; the reported medians are not themselves established upper bounds. *n*, number of experiments.

| **Carrier family** | *n* | **Papers** | **Upper-bound cases** | **Median *t*<sub>50</sub> (range), days** | **Source** |
|---|---|---|---|---|---|
| Polymeric scaffold | 6 | 4 | 5 | 0.89<sup>†</sup> (0.08–5.8) | [5,17–19] |
| Gelatin hydrogel | 1 | 1 | 0 | 1.2 | [20] |
| Composite implant | 5 | 1 | 0 | 3.9 (2.9–17.2) | [12] |
| Silk fibroin | 1 | 1 | 1 | 4.7<sup>†</sup> | [21] |
| Degradable hydrogel | 2 | 1 | 0 | 5.1 (4.2–6.0) | [22] |
| Surface coating | 2 | 1 | 0 | 5.4 (3.0–7.8) | [16] |
| Heparin affinity | 1 | 1 | 0 | 5.7 | [5] |
| ECM-coated ceramic | 2 | 1 | 0 | 7.2 (2.1–12.3) | [15] |
| Gelatin microparticle | 4 | 1 | 0 | 17.6 (3.0–20.2) | [19] |
| Ceramic or mineral | 4 | 2 | 3 | 28<sup>†</sup> (1.3–28) | [15,23] |

The dataset additionally contained 132 biological outcome records (Supplementary Table 5), 128 linked to release experiments. Linked records spanned 20 endpoint categories, including alkaline phosphatase activity and micro-computed-tomography bone volume. Stratification by endpoint, assay or animal model, time point, and anatomical site yielded 52 combinations. Fifteen combinations (29%) contained one record, and the largest contained 10. Given this heterogeneity, outcomes were summarized descriptively without pooling effect sizes.

**Table 5.**
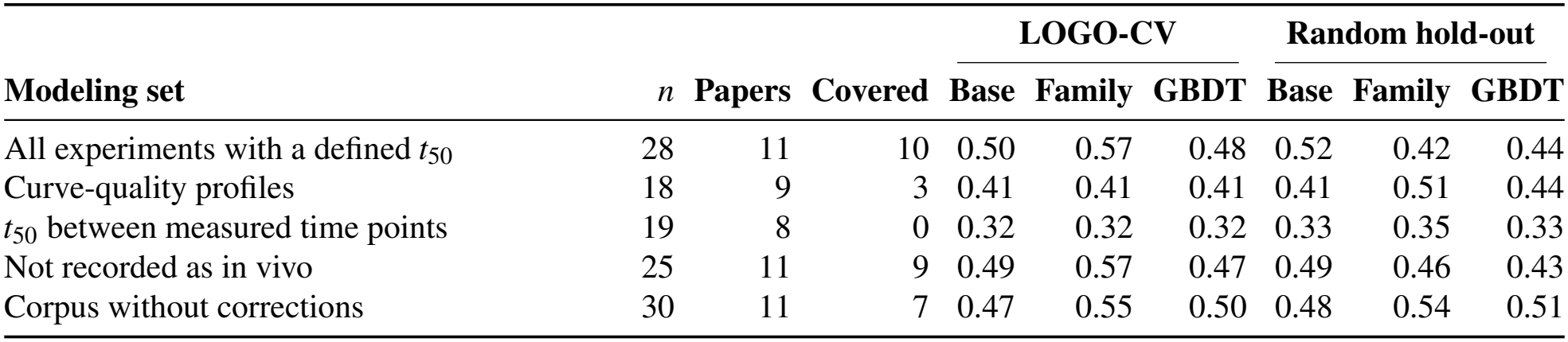
Sensitivity analysis. MAE of log_10_ *t*_50_ for the global-median baseline (Base), the family-median model (Family), and the gradient-boosted decision trees (GBDT) in leave-one-group-out cross-validation by publication (LOGO-CV) and in repeated random hold-out validation. *n*, number of experiments; Covered, experiments whose carrier family was present in the training set in LOGO-CV.

|  |  |  |  | **LOGO-CV** |  |  | **Random hold-out** |  |  |
|---|---|---|---|---|---|---|---|---|---|
| **Modeling set** | *n* | **Papers** | **Covered** | **Base** | **Family** | **GBDT** | **Base** | **Family** | **GBDT** |
| All experiments with a defined *t*<sub>50</sub> | 28 | 11 | 10 | 0.50 | 0.57 | 0.48 | 0.52 | 0.42 | 0.44 |
| Curve-quality profiles | 18 | 9 | 3 | 0.41 | 0.41 | 0.41 | 0.41 | 0.51 | 0.44 |
| *t*<sub>50</sub> between measured time points | 19 | 8 | 0 | 0.32 | 0.32 | 0.32 | 0.33 | 0.35 | 0.33 |
| Not recorded as in vivo | 25 | 11 | 9 | 0.49 | 0.57 | 0.47 | 0.49 | 0.46 | 0.43 |
| Corpus without corrections | 30 | 11 | 7 | 0.47 | 0.55 | 0.50 | 0.48 | 0.54 | 0.51 |

### 3.2 Accuracy of Values Obtained from Figures

The source-data check compared 30 image-derived values from five series in two panels of one published figure [12] with the corresponding publisher data (Figure 2; Supplementary Table 6). Day-0 values were excluded because all markers overlapped at the origin. MAE was 60.8 ng, equivalent to 1.72% of the *y*-axis range; the maximum error was 247.7 ng (4.95%). Series-specific MAEs ranged from 0.06% to 3.84% of the axis range. Twenty of 30 estimates were below the source values, suggesting a tendency toward underestimation in this comparison.

**Figure 2.**
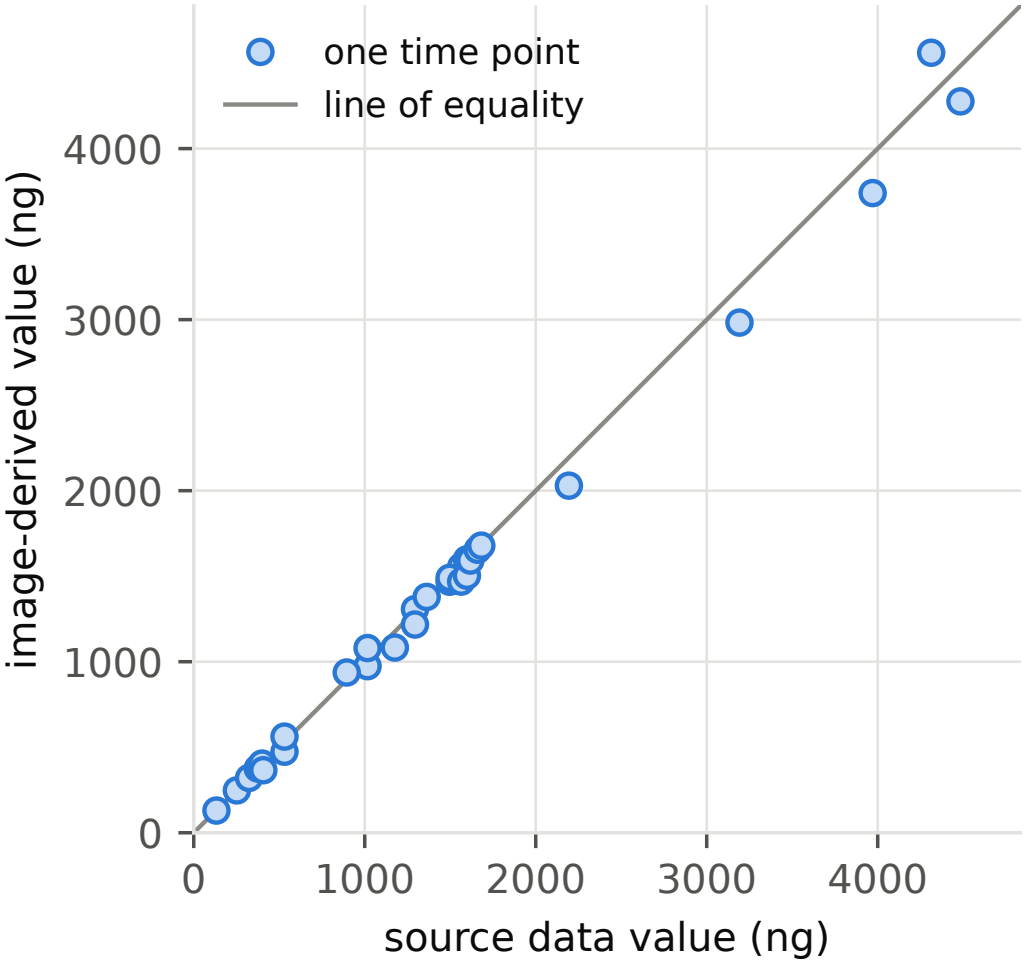
Image-derived release values compared with publisher source data for 30 time points from one paper [12]. The mean absolute error was 60.8 ng (1.72% of the *y*-axis range).

Six profiles were selected for the second-pass check (Supplementary Table 7). Of their points, 27 manually digitized values across four profiles were read a second time. Readings differed by 0.11 percentage points on average and at most 1 percentage point. The remaining 21 points were numerical source data (7 points), text-reported or text-derived values (9), or calculations from published phase rates (5), and were not redigitized.

These checks provide limited validation. The source-data comparison covers script-based extraction from one clear figure, whereas the second-pass check assesses repeatability across four manually read profiles. Neither establishes the accuracy of all manually digitized curves. Provenance labels were therefore retained throughout the analyses.

### 3.3 Comparability of the Release Records

Four dimensions constrain the comparability of release records (Figure 3).

**Figure 3.**
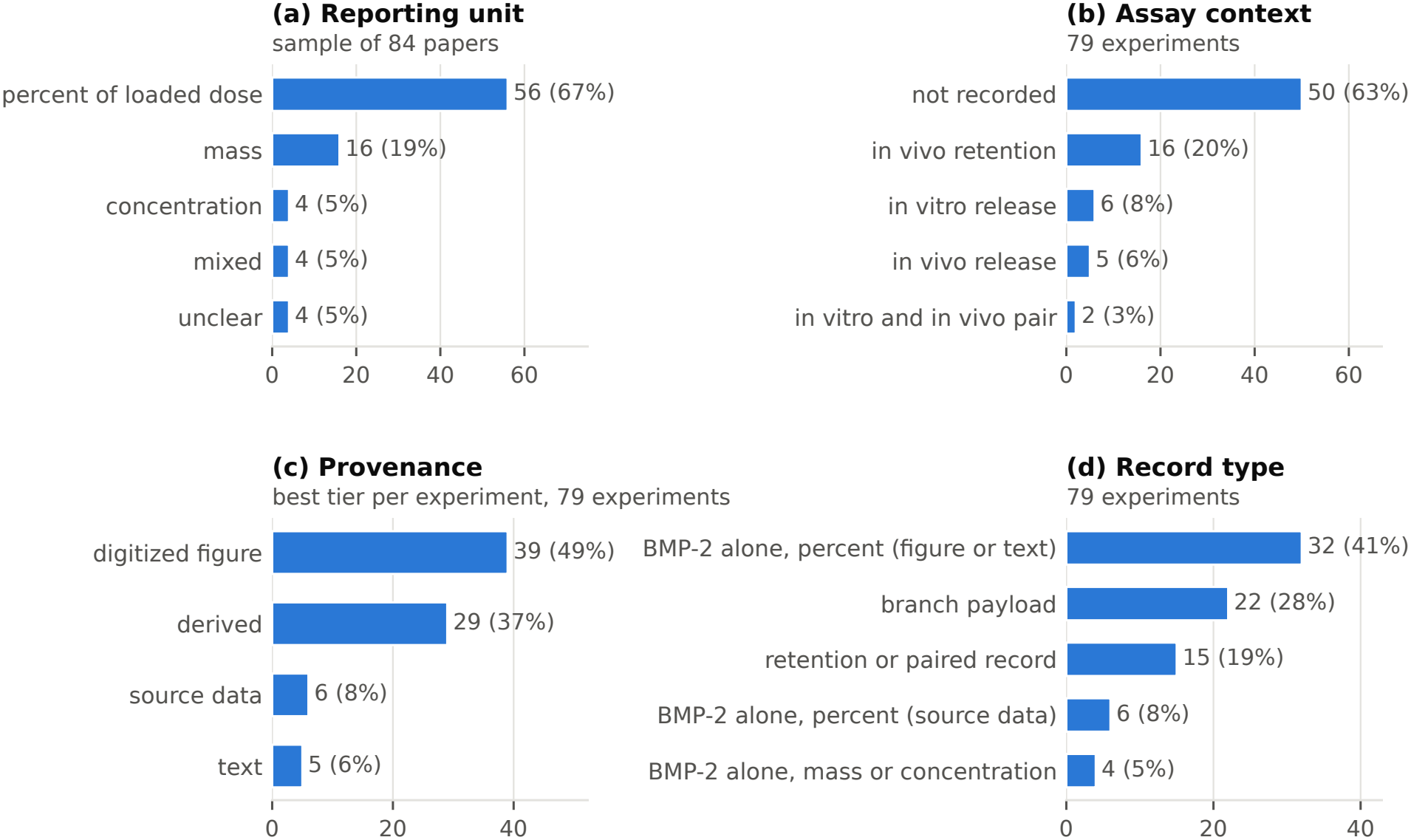
Comparability of the release records. (**a**) Reporting unit in a sample of 84 papers. (**b**) Assay context of the 79 experiments. (**c**) Best provenance level of each experiment. (**d**) Record type of each experiment; the two groups with BMP-2 alone in percent form the percent-release cohorts. Percentages refer to the total of each panel.

#### Reporting unit

In the separate 84-paper sample (Supplementary Table 8), 56 papers (67%) reported release as a percentage of the loaded dose, 16 (19%) as mass, and 4 (5%) as concentration. Four (5%) used mixed units and 4 (5%) had unclear units. Thus, 28 papers (33%) did not report release exclusively in percentage units. These classifications were not manually verified.

#### Assay context

Context was unrecorded for 50 of 79 experiments (63%). Sixteen (20%) concerned in vivo retention, 6 (8%) in vitro release, 5 (6%) in vivo release, and 2 (3%) paired in vitro/in vivo measurements. Release into buffer and retention at an implant site measure different processes; in vitro BMP-2 profiles do not reliably predict in vivo release across formulations [14].

#### Provenance

Classification by each experiment’s highest available provenance assigned 39 experiments (49%) to digitized figures, 29 (37%) to derived values, 6 (8%) to publisher source data, and 5 (6%) to text.

#### Record type

The percent-release cohorts contained 38 experiments (48%): 6 with source data and 32 with figure-derived, text-derived, or calculated measurements. The remaining records comprised 22 branch-payload experiments (28%), 15 retention or paired in vitro/in vivo records (19%), and 4 mass-or concentration-only experiments (5%).

Release-rate analyses (Sections 3.4 and 3.5) were restricted to the 38 percent-release experiments. Al-though these share payload and reporting scale, assay context was unrecorded for 29; 6 were in vitro and 3 were in vivo.

### 3.4 Release Rate by Carrier Family

Twenty-eight of the 38 percent-release experiments had a defined *t*_50_, representing 11 publications and 10 carrier families (Table 4; Figure 4). Nineteen estimates were bracketed by measured time points and ranged from 1.2 to 20.2 days. The other 9 were upper-bound cases: release already exceeded 50% at the first observation, occurring at 4 hours (2 experiments), 1 day (3), 7 days (1), or 28 days (3). Final cumulative release ranged from 51% to 100%, with final sampling times between 7 and 35 days.

**Figure 4.**
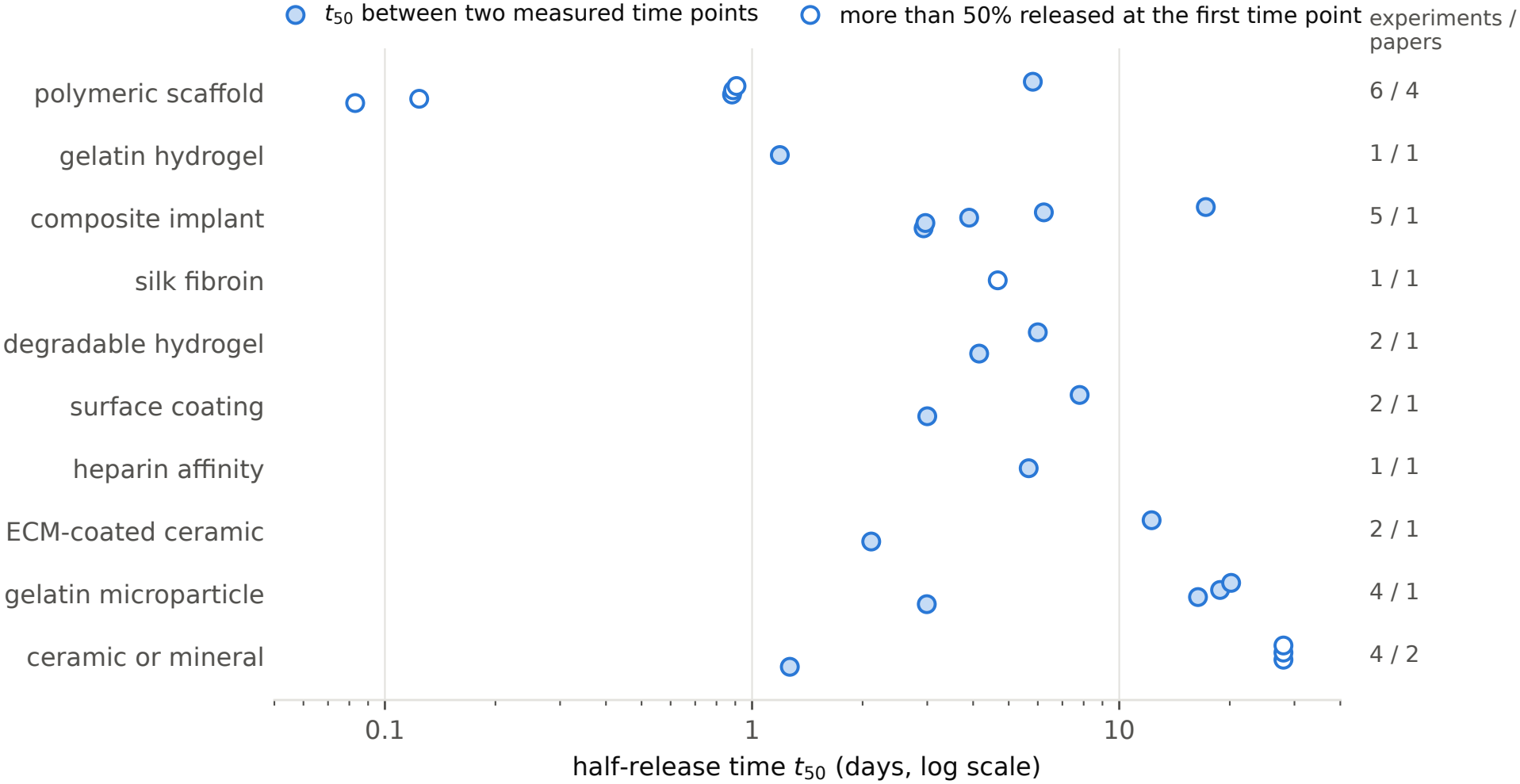
Half-release time (*t*_50_) of the 28 experiments in the modeling set, by carrier family. Families are ordered by their median. Open markers identify upper-bound cases, in which release exceeded 50% at the first observation; plotted values follow the estimation convention in Section 2.6. The numbers on the right give the experiments and papers in each family.

Family medians had limited independent support. Eight of ten families, accounting for 18 of 28 experiments, were represented by a single publication; three families contained only one experiment. Only polymeric scaffolds (four publications) and ceramics or minerals (two publications) had multi-publication coverage, and most *t*_50_ records in these families were upper-bound cases.

Within-publication variation was substantial. Seven publications contributed multiple experiments; one reported only a single time point, precluding comparison of release kinetics. In each of the remaining six, the slowest release occurred with an added binding component, coating layer, or higher crosslinking. For example, heparin-conjugated poly(lactide-co-glycolide) had a *t*_50_ of 5.7 days, whereas the unmodified polymer released its entire dose by the first observation at 4 hours [5]. For *β* -tricalcium phosphate, *t*_50_ increased from 1.3 days without a coating to 2.1 and 12.3 days with cell-derived extracellular matrix (ECM) coatings [15]. Chitosan-coated titanium yielded 7.8 days, compared with 3.0 days for BMP-2 on titanium [16]. An additional polymer layer increased implant *t*_50_ from 3.9 to 17.2 days [12]. Across these six publications, within-study estimates varied by factors of 1.4 to more than 30, consistent with prolonged retention through protein binding and additional diffusion barriers [4–6].

### 3.5 Prediction of the Half-Release Time from Carrier Family

#### Association in the full modeling set

Carrier family accounted for 57% of the variance in log_10_ *t*_50_ (*η*^2^ = 0.57), decreasing to 35% after adjustment for group count (*ε*^2^ = 0.35; permutation *p* = 0.05). For publication, *η*^2^ = 0.53 and *ε*^2^ = 0.25 (*p* = 0.13). Carrier family and publication were strongly associated (bias-corrected Cramér’s *V* = 0.80), with eight families represented by one publication each. Family effects could therefore not be reliably separated from publication-specific differences in laboratory procedures, assay conditions, or dose.

#### Random hold-out validation

MAE was 0.52 *±* 0.15 for the global-median baseline, 0.42 *±* 0.14 for the family-median model, and 0.44 *±* 0.13 for GBDT (mean *±* standard deviation across 200 repetitions; Figure 5a). The family-median and GBDT models outperformed the baseline in 141 and 146 repetitions, respectively. However, 81% of test experiments shared a publication with training experiments; 14% belonged to families absent from training (Figure 5b).

**Figure 5.**
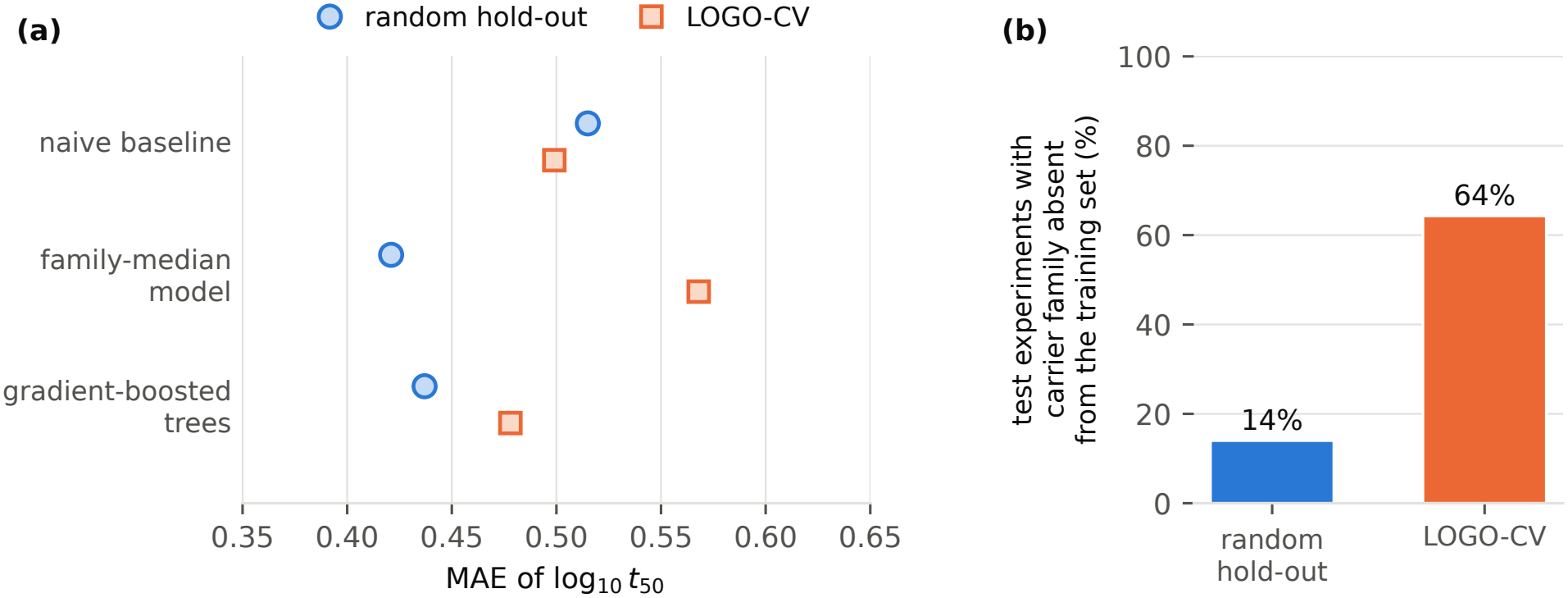
Prediction of log_10_ *t*_50_ from the carrier family in the modeling set (28 experiments). The global-median baseline is labeled “naive baseline” in the figure. (**a**) Mean absolute error (MAE) of the three models in repeated random hold-out validation and in leave-one-group-out cross-validation by publication (LOGO-CV). (**b**) Test experiments whose carrier family was absent from the training set.

#### Publication-grouped cross-validation

LOGO-CV MAE was 0.50 for the baseline, 0.57 for the family-median model, and 0.48 for GBDT. Relative to the baseline, differences were 0.07 (95% publication-bootstrap interval, *−* 0.08 to 0.22) and *−* 0.02 (*−* 0.14 to 0.14), respectively. Thus, publication-grouped validation did not establish an improvement from either family-based model.

Eighteen of 28 held-out experiments (64%) belonged to families absent from training. For these records, the family-median model reverted to the baseline; GBDT predicted from the remaining encoded categories without training examples of the held-out family. MAE was 0.29 for both the baseline and family-median model and 0.30 for GBDT. The remaining 10 experiments, from the polymeric scaffold and ceramic or mineral families, had training-set family coverage. Their MAEs were 0.87, 1.07, and 0.81, respectively; 8 of these 10 were upper-bound cases.

Evaluating a family’s transfer across publications requires representation in at least two independent publications. Eight of ten families failed this condition, each requiring at least one additional independent study to permit such evaluation.

#### Sensitivity analysis

Performance depended on modeling-set composition (Table 5). Across all LOGO-CV analyses, family-based MAEs ranged from 0.02 below to 0.08 above the baseline. Under random hold-out validation, both family-based models improved on the baseline in the full set and the subset excluding records identified as in vivo. For curve-quality profiles, profiles with bracketed *t*_50_, and the uncorrected corpus, their MAEs were similar to or higher than the baseline. In the bracketed-*t*_50_ subset, each family occurred in only one publication; no held-out family was represented in training, and all three LOGO-CV MAEs were 0.32.

## 4 Discussion

This exploratory analysis examined BMP-2 release records from 21 publications using a minimal prediction task. Four findings characterize the available evidence. First, approximately half of the experiments shared BMP-2-only payloads and a percentage-of-loaded-dose scale. Second, script-based readings from one clear figure differed from numerical source data by 1.7% of the axis range on average and at most 5.0%. Third, within-study comparisons reproduced release-rate patterns expected from established binding and barrier mechanisms. Fourth, the apparent benefit of carrier family under random splitting was not established when entire publications were held out.

The validation discrepancy is consistent with a predictor that is strongly associated with data source. Under random splitting, most test experiments shared a publication with training experiments. Carrier family can then act as a proxy for publication, and measured performance primarily reflects prediction of additional formulations from familiar studies. It does not establish transfer to a new study. Grouping by drug–polymer combination served a related purpose in Bannigan et al. [7]. The present findings support publication-grouped validation for literature-derived release datasets.

The LOGO-CV results nevertheless remain inconclusive. Only 10 experiments had families represented in training, and 8 were upper-bound cases. Bootstrap intervals for model–baseline differences were wide and included zero. Consequently, these data cannot establish whether carrier family predicts half-release time in an independent study.

Three dataset properties constrain interpretation. First, predictor coverage was limited to one categorical variable with 10 levels across 28 experiments. By comparison, Bannigan et al. used 181 profiles and 17 features describing drug, polymer, formulation, time, and early release [7]. Broad family labels conceal variation in composition, crosslinking, dose, and assay conditions; within gelatin microparticles alone, *t*_50_ ranged from 3.0 to 20.2 days. Second, strong family–publication association prevents reliable separation of material effects from study effects. Laboratories typically vary formulations within a single carrier system, while controlled cross-family comparisons are uncommon. Third, the small modeling set was sensitive to removing or relabeling a few records (Table 5). The random-split advantage was not reproduced after excluding upper-bound cases or retaining the original labels for the three polymer-scaffold controls.

Observed release patterns align with established BMP-2 carrier behavior. Heparin-containing materials exploit the protein’s heparin-binding site [4] to prolong retention [5,6], whereas collagen sponges exhibit rapid initial release [1,2]. Within-study comparisons similarly associated binding components, coating layers, and increased crosslinking with slower release (Section 3.4). Reproducing these expected patterns provides a consistency check on extraction, although it does not independently validate every record. Assay medium also matters: serum proteins competed with BMP-2 for heparin binding in one included study [6], reinforcing the need to capture assay conditions explicitly.

Three diagnostic checks should precede model fitting on literature-derived datasets. First, quantify associations between predictors and publication, using statistics such as bias-corrected Cramér’s *V* for categorical variables. Second, compare publication-grouped validation with random splitting to distinguish within-study performance from cross-study transfer. Third, report training-set coverage of held-out feature values, particularly categories that have no examples in training. These checks are computationally inexpensive and expose limitations that aggregate error estimates can conceal.

Data curation benefits from value-level provenance and source references, deterministic normalization rules, human verification, and documented decisions. Checks must address record identity and interpretation as well as numerical transcription. In this compilation, removing two duplicate records and correcting three carrier-family labels changed the prediction results (Section 2.5; Table 5). This sensitivity illustrates why seemingly minor classification decisions can materially affect results in a small literature-derived sample.

Four improvements would strengthen predictive modeling. First, each family requires representation in multiple independent publications; only two of ten families currently meet the minimum of two. Second, quantitative formulation and assay descriptors should be structured as predictors, including BMP-2 dose, loading method, geometry, crosslink density, release medium, and assay context. These variables were not available as structured predictors in the reported analysis.

Third, numerical release data should accompany publications. Such data were available for only one of the 21 contributing papers. Earlier sampling is also necessary to resolve *t*_50_ when release exceeds 50% before the first observation. Fourth, prospective experiments should vary defined material properties while controlling other conditions. For example, a graded heparin-density hydrogel series could use a fixed BMP-2 dose, geometry, medium, and sampling schedule. Release should be followed to a plateau, with residual carrier-associated protein quantified at the endpoint.

The convenience sample was shaped by the source spreadsheet and access to full texts and figures; reported proportions need not represent the complete literature. The modeling set contained only 28 experiments from 11 publications, including 9 upper-bound cases. Models used one categorical predictor, and GBDT hyperparameters were not tuned. The findings therefore characterize this dataset rather than establish a general inability of carrier properties to predict release.

Assay context was unrecorded for 63% of experiments. The modeling set mixed in vitro and in vivo records, and excluding explicitly identified in vivo records could not resolve uncertainty in the unclassified experiments. Carrier labels lacked independent second-reader assessment, and three required correction. Manual digitization used visual estimates without dedicated software; repeatability was assessed in four profiles by non-independent rereading, while source-data validation covered a single figure. Candidate proposal acceptance was not logged, preventing quantification of the LLM assistant’s contribution. Reporting-unit labels for the separate 84-paper sample were model-assigned without human verification. Finally, final release depended on observation duration, which ranged from 7 to 35 days, and outcome heterogeneity precluded pooled biological-effect analysis.

## 5 Conclusions

This exploratory analysis summarizes 79 BMP-2 release experiments and 487 time points from 21 publications. Approximately half the experiments share a BMP-2-only payload and percentage-of-loaded-dose scale. Within-study comparisons reproduce expected effects of binding components and coatings, but apparent predictive gains from carrier family under random splitting were not established under publication-grouped validation. Strong family–publication association, limited training coverage of held-out families, and sensitivity to a few record corrections constrain cross-study inference. More independent studies per family, structured quantitative carrier and assay descriptors, and accessible numerical source data are needed to evaluate generalizable BMP-2 release models. The present findings motivate these requirements rather than establish a deployable release-prediction model.

## Supplementary Materials

The following supporting information is provided with this article. Supplementary Table 1: the 30 publications in the corpus. Supplementary Table 2: the 79 release experiments with labels and release metrics. Supplementary Table 3: the two removed duplicate records and the three corrected carrier-family labels. Supplementary Table 4: the 487 release time points with provenance and source reference. Supplementary Table 5: the 132 biological outcome records. Supplementary Table 6: image-derived values compared with publisher source data. Supplementary Table 7: second-pass check of manually digitized points. Supplementary Table 8: reporting units in the separate 84-paper sample. Supplementary Data 1: the source spreadsheet of 1051 publications on BMP-2 delivery. Supplementary Data 2: analysis scripts, input tables, and result files.

## Author Contributions

W.Z.: conceptualization, methodology, data curation, formal analysis, investigation, visualization, writing of the original draft; L.Z.: data curation, investigation, validation; W.H.: validation, review and editing of the manuscript; Y.P.Y.: supervision, review and editing of the manuscript. All authors have read and agreed to the published version of the manuscript.

## Funding

This research was funded by a Stanford Bio-X Seed Grant.

## Institutional Review Board Statement

Not applicable. This study analyzed previously published data and did not involve humans or animals.

## Informed Consent Statement

Not applicable.

## Data Availability Statement

Aggregate results are reported in the main-text tables and figures. The record-level extraction tables, analysis scripts, and result files are provided in the Supplementary Materials. No new experimental data were generated.

## Conflicts of Interest

The authors declare no conflict of interest.

## Use of AI and AI-assisted Technologies

During the preparation of this work, the authors used large language model assistants (Claude, Anthropic; Codex, OpenAI) to support literature screening and candidate data extraction as described in the Methods, and to assist with analysis code and with drafting and editing the text. After using these tools, the authors reviewed and edited the content as needed and take full responsibility for the content of the published article.

## Notes

### Competing Interest Statement

The authors have declared no competing interest.

## References

[1] Friess, W.; Uludag, H.; Foskett, S.; Biron, R.; Sargeant, C. Characterization of absorbable collagen sponges as rhBMP-2 carriers. Int. J. Pharm. 1999, 187, 91–99. 10.1016/S0378-5173(99)00174-X

[2] Jeon, S.-O.; Lamichhane, S.; Oh, D.-H.; et al. Engineering a BMP2–risedronate complex with sustained release for osteoporosis therapy. Arch. Pharm. Res. 2025, 48, 1001–1021. 10.1007/s12272-025-01568-8

[3] Carragee, E.J.; Hurwitz, E.L.; Weiner, B.K. A critical review of recombinant human bone morphogenetic protein-2 trials in spinal surgery: Emerging safety concerns and lessons learned. Spine J. 2011, 11, 471–491. 10.1016/j.spinee.2011.04.023

[4] Ruppert, R.; Hoffmann, E.; Sebald, W. Human bone morphogenetic protein 2 contains a heparin-binding site which modifies its biological activity. Eur. J. Biochem. 1996, 237, 295–302. 10.1111/j.1432-1033.1996.0295n.x

[5] Jeon, O.; Song, S.J.; Kang, S.-W.; Putnam, A.J.; Kim, B.-S. Enhancement of ectopic bone formation by bone morphogenetic protein-2 released from a heparin-conjugated poly(L-lactic-co-glycolic acid) scaffold. Biomaterials 2007, 28, 2763–2771. 10.1016/j.biomaterials.2007.02.023

[6] Hettiaratchi, M.H.; Chou, C.; Servies, N.; Smeekens, J.M.; Cheng, A.; Esancy, C.; Wu, R.; McDevitt, T.C.; Guldberg, R.E.; Krishnan, L. Competitive protein binding influences heparin-based modulation of spatial growth factor delivery for bone regeneration. Tissue Eng. Part A 2017, 23, 683–695. 10.1089/ten.tea.2016.0507

[7] Bannigan, P.; Bao, Z.; Hickman, R.J.; Aldeghi, M.; Häse, F.; Aspuru-Guzik, A.; Allen, C. Machine learning models to accelerate the design of polymeric long-acting injectables. Nat. Commun. 2023, 14, 35. 10.1038/s41467-022-35343-w

[8] Bao, Z.; Kim, J.; Kwok, C.; Le Devedec, F.; Allen, C. A dataset on formulation parameters and characteristics of drug-loaded PLGA microparticles. Sci. Data 2025, 12, 364. 10.1038/s41597-025-04621-9

[9] He, S.; Barón, A.; Munteanu, C.R.; de Bilbao, B.; Casañola-Martin, G.M.; Chelu, M.; Musuc, A.M.; Bediaga, H.; Ascencio, E.; Castellanos-Rubio, I.; Arrasate, S.; Pazos, A.; Insausti, M.; Rasulev, B.; González-Díaz, H. Drug release nanoparticle system design: Data set compilation and machine learning modeling. ACS Appl. Mater. Interfaces 2025, 17, 5290–5306. 10.1021/acsami.4c16800

[10] Gartlehner, G.; Kugley, S.; Crotty, K.; Viswanathan, M.; Dobrescu, A.; Nussbaumer-Streit, B.; Booth, G.; Tread-well, J.R.; Han, J.M.; Wagner, J.; Apaydin, E.A.; Coppola, E.L.; Maglione, M.; Hilscher, R.; Chew, R.; Pilar, M.; Swanton, B.; Kahwati, L.C. Artificial intelligence-assisted data extraction with a large language model: A study within reviews. Ann. Intern. Med. 2025, 178, 1763–1771. 10.7326/ANNALS-25-00739

[11] Khraisha, Q.; Put, S.; Kappenberg, J.; Warraitch, A.; Hadfield, K. Can large language models replace humans in systematic reviews? Evaluating GPT-4’s efficacy in screening and extracting data from peer-reviewed and grey literature in multiple languages. Res. Synth. Methods 2024, 15, 616–626. 10.1002/jrsm.1715

[12] Lin, S.; Maekawa, H.; Moeinzadeh, S.; Lui, E.; Alizadeh, H.V.; Li, J.; Kim, S.; Poland, M.; Gadomski, B.C.; Easley, J.T.; Young, J.; Gardner, M.; Mohler, D.; Maloney, W.J.; Yang, Y.P. An osteoinductive and biodegradable intramedullary implant accelerates bone healing and mitigates complications of bone transport in male rats. Nat. Commun. 2023, 14, 4455. 10.1038/s41467-023-40149-5

[13] Bergsma, W. A bias-correction for Cramér’s V and Tschuprow’s T. J. Korean Stat. Soc. 2013, 42, 323–328. 10.1016/j.jkss.2012.10.002

[14] Olthof, M.G.L.; Tryfonidou, M.A.; Dadsetan, M.; Dhert, W.J.A.; Yaszemski, M.J.; Kempen, D.H.R.; Lu, L. In vitro and in vivo correlation of BMP-2 release profiles from complex delivery vehicles. Tissue Eng. Part C Methods 2018, 24, 379–390. 10.1089/ten.TEC.2018.0024

[15] Kang, Y.; Kim, S.; Khademhosseini, A.; Yang, Y. Creation of bony microenvironment with CaP and cell-derived ECM to enhance human bone-marrow MSC behavior and delivery of BMP-2. Biomaterials 2011, 32, 6119–6130. 10.1016/j.biomaterials.2011.05.015

[16] Han, L.; Lin, H.; Lu, X.; Zhi, W.; Wang, K.-F.; Meng, F.-Z.; Jiang, O. BMP2-encapsulated chitosan coatings on functionalized Ti surfaces and their performance in vitro and in vivo. Mater. Sci. Eng. C 2014, 40, 1–8. 10.1016/j.msec.2014.03.043

[17] Srouji, S.; Ben-David, D.; Lotan, R.; Livne, E.; Avrahami, R.; Zussman, E. Slow-release human recombinant bone morphogenetic protein-2 embedded within electrospun scaffolds for regeneration of bone defect: In vitro and in vivo evaluation. Tissue Eng. Part A 2011, 17, 269–277. 10.1089/ten.tea.2010.0250

[18] Kim, B.-R.; Nguyen, T.B.L.; Min, Y.-K.; Lee, B.-T. In vitro and in vivo studies of BMP-2-loaded PCL–gelatin–BCP electrospun scaffolds. Tissue Eng. Part A 2014, 20, 3279–3289. 10.1089/ten.tea.2014.0081

[19] Patel, Z.S.; Yamamoto, M.; Ueda, H.; Tabata, Y.; Mikos, A.G. Biodegradable gelatin microparticles as delivery systems for the controlled release of bone morphogenetic protein-2. Acta Biomater. 2008, 4, 1126–1138. 10.1016/j.actbio.2008.04.002

[20] Chen, F.; Wu, Z.; Wang, Q.; Wu, H.; Zhang, Y.; Nie, X.; Jin, Y. Preparation and biological characteristics of recombinant human bone morphogenetic protein-2-loaded dextran-co-gelatin hydrogel microspheres, in vitro and in vivo studies. Pharmacology 2005, 75, 133–144. 10.1159/000088212

[21] Karageorgiou, V.; Tomkins, M.; Fajardo, R.; Meinel, L.; Snyder, B.; Wade, K.; Chen, J.; Vunjak-Novakovic, G.; Kaplan, D.L. Porous silk fibroin 3-D scaffolds for delivery of bone morphogenetic protein-2 in vitro and in vivo. J. Biomed. Mater. Res. Part A 2006, 78A, 324–334. 10.1002/jbm.a.30728

[22] Holloway, J.L.; Ma, H.; Rai, R.; Burdick, J.A. Modulating hydrogel crosslink density and degradation to control bone morphogenetic protein delivery and in vivo bone formation. J. Control. Release 2014, 191, 63–70. 10.1016/j.jconrel.2014.05.053

[23] Zhao, J.; Wang, S.; Bao, J.; Sun, X.; Zhang, X.; Zhang, X.; Ye, D.; Wei, J.; Liu, C.; Jiang, X.; Shen, G.; Zhang, Z. Trehalose maintains bioactivity and promotes sustained release of BMP-2 from lyophilized CDHA scaffolds for enhanced osteogenesis in vitro and in vivo. PLoS ONE 2013, 8, e54645. 10.1371/journal.pone.0054645

